# Personalized Morphology, Replication Timing, and RNA based Gene Expression Networks for Basal-like and Classical subtyping genes in Pancreatic Adenocarcinoma

**DOI:** 10.64898/2026.03.12.711434

**Authors:** Alejandro Leyva, M. Khalid Khan Niazi

## Abstract

Network biology traditionally identifies gene correlations that reflect biological pathways. While LIONESS enables individualized gene networks, the influence of replication timing on these correlations remains unexplored. Replication timing reflects the temporal order of DNA synthesis and is tightly linked to chromatin state, methylation, and transcriptional stability, all of which affect tumor behavior. Integrating replication-timing proxies derived from methylation data therefore offers a bridge between epigenetic state and functional gene coordination, while morphology provides an additional route for inferring gene expression.

This is the first study to integrate replication-timing proxies and morphological embeddings into individualized LIONESS gene networks. The aim is to determine how replication timing and morphology derived from bulk methylation and image embeddings influence gene coexpression in pancreatic cancer. Patient-specific networks were generated for basal and classical pancreatic ductal adenocarcinoma subtypes using TCGA data.

Results show an 80% AUC for RNA-replication-timing–based subtype prediction modules and a 75% AUC for morphology-based networks. Incorporating replication timing and morphology increased network robustness while maintaining classification performance. Notably, the 80% AUC was achieved using only 17 of the 50 Moffitt genes, with 16 overlapping the PURIST gene set, indicating that replication timing captures clinically relevant regulatory structure. These findings suggest that replication-timing proxies can act as epigenetic indicators of mechanistic gene coordination and may help identify patients with distinct replication stress or chromatin accessibility profiles relevant to therapeutic response.

## 1 Introduction

Genes act in cascades that are regulated by intermediary messengers, such that inhibitory gene expression is inversely correlated with exhibitory gene expression. Genes that are correlated in cascades can reveal mechanistic correlations between genes involved in pathways outside of immediate functional domains. Network biology provides a means to study gene coexpression by building networks of genes corresponding to pairwise associations, with edges defined by correlation strength [10,11].

Replication timing refers to the temporal order in which genomic regions are duplicated during S phase and is closely tied to chromatin stability, accessibility, proliferation, and transcriptional potential [2,18]. Replication timing is often measured using Repli-seq, a method in which bromodeoxyuridine is incorporated into replicating DNA, enabling immunoprecipitation and quantification of newly synthesized sequences [1]. Replication timing domains are positively correlated with methylation when log-scaled, but methylation is inversely correlated with replication timing on normalized scales, indicating that highly methylated regions replicate later [5]. Because methylation correlates with replication lateness, an inverted and smoothed methylation signal can serve as a replication-timing proxy [4].

In cancer, elevated replication timing scores correlate with increased gene expression and oncogenic activity [17]. Regions harboring oncogenes such as MYC replicate earlier, reinforcing aberrant transcription in tumor cell lines [1,3]. DNA damage and repair genes also show replication-timing–dependent responsiveness to standard-of-care chemotherapies such as FOLFIRINOX [17]. Consequently, replication timing offers a mechanistic basis for modeling differences in cascade activity and gene coexpression with respect to chromatin structure.

Pancreatic cancer includes two major transcriptomic subtypes, Basal-Like and Classical, defined by the expression of 50 genes identified by Moffitt et al. [15]. These subtypes correlate with morphological phenotypes and contain anticorrelated gene sets. Basal-Like tumors are poorly differentiated and progenitor-like, with worse prognosis and weaker response to chemotherapy, while Classical tumors exhibit epithelial features and improved treatment response [15]. Clinically, PURIST subtyping refines this classification using 16 of the strongest predictor genes, assessed pairwise rather than by ranked expression [16].

Gene expression has been modeled extensively in pancreatic cancer using RNA-seq, but modeling genetic coexpression structure—how genes interact—remains underexplored. LIONESS networks enable personalized gene coexpression analysis at the patient level [7]. Although prior studies have modeled replication timing and gene expression jointly [2,6,17], none have done so at single-patient resolution. Moreover, no prior work integrates morphological features extracted from whole-slide images using deep learning. By incorporating vision-transformer embeddings, genetic coexpression can be analyzed alongside physical tissue characteristics.

While RNA sequencing supports coexpression network construction, very few studies integrate replication-timing signals into multimodal networks, and none have developed individualized patient-level multimodal LIONESS networks. LIONESS supports the construction of such networks and enables mechanistic interpretation of coexpression patterns in patients [7]. Understanding these relationships allows inference of downstream gene activity within shared cascades.

Pancreatic ductal adenocarcinoma is the tenth most diagnosed cancer in the United States, yet the third leading cause of cancer-related deaths. It is diagnosed late and is highly aggressive. Transcriptomic subtyping has been used to identify patients with poor prognosis, treatment resistance, and distinct cellular differentiation patterns [15,16]. While the gene cascades underlying these subtypes are known, replication-timing contributions to subtype-specific coexpression remain unexplored. Graph-based networks allow replication-timing proxies to influence node-to-node relationships and reveal mechanistic structure in coexpression topology.

The objective of this study is to determine how replication timing and morphology influence gene coexpression structure in pancreatic cancer. We integrate replication-timing proxies, vision-transformer embeddings, and patient-level LIONESS networks to evaluate their contributions to subtype prediction and mechanistic network stability.

## 2 Materials and Methods

A total of 183 patients from TCGA PAAD were genetically subtyped using Moffitt-Fifty-gene subtyping was mapped to methylation proxies and embeddings at a 1:1 case–embedding ratio [15]. Methylation was preprocessed using SeSAMe and indexed using the Illumina 450K assay [5]. Raw TXT methylation files were parsed using an automatic column-detection pipeline that identifies probe IDs and beta values across heterogeneous vendor formats. Beta values were coerced to numeric form, clipped to the range 0 to 1, and matched to Illumina manifest probe identifiers after normalization of probe strings. Promoter-associated probes were selected using canonical Illumina annotations (TSS200, TSS1500, 5UTR, First Exon), and probes mapping to multiple genes were expanded so that each gene received its corresponding probe values. For each sample, promoter beta values were averaged across all probes assigned to each gene to produce a reproducible promoter-level methylation estimate [5]. Gene-wise z-scores were computed across samples to normalize differences in probe density and distribution. Methylation states were summarized as hypo, baseline, or hyper using fixed z-score thresholds. Both long (sample × gene) and wide (gene × sample) methylation matrices were generated for downstream network analysis. This procedure ensured consistent promoter-level aggregation and reproducible methylation-derived replication-timing proxies across the cohort.

Methylation was converted into replication timing by inversion, z-scoring, and smoothing, following the established correlation between methylation and replication-timing domains [4,5].

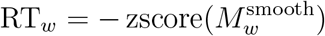

Smoothing is then applied across the domains to normalize methylation since B matrices have been preprocessed.

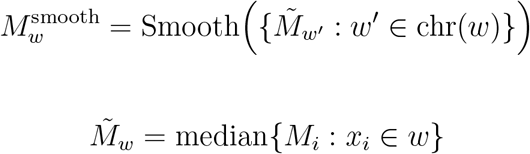

After Smoothing, zscoring is applied to the methylation domains such that higher beta values regions will have lower replication timing and vice-versa. The Replication timing for each domain is calculated using the averages of the inversed Beta values:

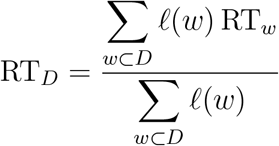

where l(w) is the genomic window length, which is weighted based on the length of each domain. if the bin length is lower, then it is calculated using the following equation:

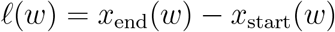

If the bins are uniform, then:

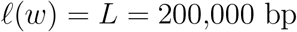

The fraction of domains that have late replication timing, as well as the sample level averaged replication timing is calculated as follows:

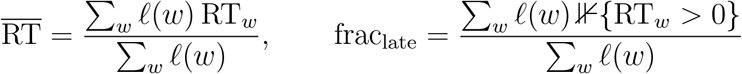

After the replication timing domains are created into 300kb regions, the networks are constructed using gene correlations, creating a global correlation network across all patients. correlation of genes within replication timing domains and the lateness of replication would influence the strength and connection between edges.

**Table 1:** Key parameters used for individualized replication–timing and RNA network generation.

| Parameter | Value |
| --- | --- |
| Absolute minimum correlation | 0.5 |
| Global top-K edges | 5000 |
| Plot top-K edges | 500 |
| Minimum non-missing samples | 5 |
| Random seed | 1337 |
| Replication-timing / RNA mix | 1.0, 0.5, 0.5 |
| Number of RT shuffles | 100 |
| Grid search range | 0.5:1.5:0.25, 0:1:0.25, 0:1:0.25 |
| Partial correlation used | 1 |
| Bootstrap iterations | 50 |
| Bootstrap fraction | 0.8 |
| Top-K for stability | 10000 |
| Replication-timing threshold | 0.0 |

The absolute minimum Pearson correlation any two expressed genes must have is 0.5 or above; otherwise, the network becomes oversaturated with edges. After calculating the number of edges, the top 5,000 weights between genes are used, and only the top 500 of those weighted edges are plotted in the network. Within a gene, a minimum of five samples is required to be considered for network parameterization. Random seeding is used to create replicable results within the dataset. The RNA-mix parameters are the scaling parameters used for the subtyping task for alpha, beta, and gamma. A total of 100 shuffles were performed, whereby the labels were permuted to create null control graphs, ensuring that variation is not due to dataset artifacts. For each sample, the parameters for alpha, beta, and gamma were modified to visualize changes in network behavior and saturation with changes to the subtyping task. Only one partial correlation matrix of replication timing and RNA was generated. Fifty iterations were used to estimate edge stability, and only 80 percent of those networks were used per bootstrap. The top 10,000 edges were used to compute Jaccard stability between the edges of each network. The replication-timing correlation was set to zero to visualize changes in network behavior with the introduction of replication timing.

As shown, gene associations via Pearson correlation were established between pairs of genes and their expression, and the optional partial correlation network was created by using the inverse covariance matrices of gene expression, labeled as omega. This, in turn, can be used to produce the partial correlation between two pairs of expressed genes, labeled as rho.

### Gene associations

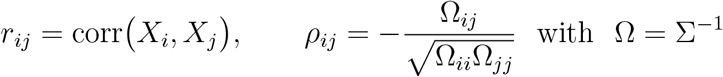

Replication timing correlations are produced by using the replication timing average of a Domain, where i and j are genes, where as u and v are all of the genes within the domains of i and j.

### Replication-timing similarity

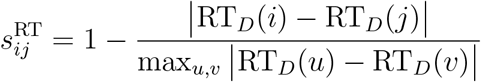

The edge weights between gene pairs are influenced by the similiarity scores between RT domains for each gene pair, the correlation between two genes, and the norm of the correlation between gene pairs using z scoring. This produces an RT influenced RNA graph, integrating replicative domains and the correlation between genes within the same replication domains.

### Integrated edge weights

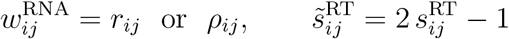

For the integrated RT_R_*NAnetwork, wecomputedforeachgenepairthePearsoncorrelationofexpressionandtheP timingprofiles*.*Replication*−*timingvalueswererank*−*transformedbeforecorrelation*.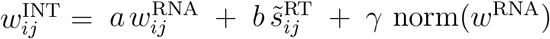 with mixing coefficients *a, b, γ* chosen by the grid ranges reported in Methods.

The edge weights are thresholded using parameter Tau reported in the methods table (r=.5). The top number of edge weights specified in the parameter table are then used to create and construct the global network. Patient-specific networks were estimated with LIONESS using the RT_R_*NAintegratedsimilarityfunction*; *RNA*−*onlyandRT* −*onlynetworkswereanalyzedatthegloballevelonly.*

### Edge selection

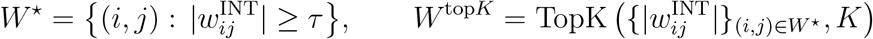

LIONESS networks were constructed to create personalized networks for each patient. Edges are computed using the difference between any edge across the global network, and subtracting the inputs without the specific sample, whereby that discrepancy changes the edge weights in the patient network.

### Individualized networks (LIONESS)

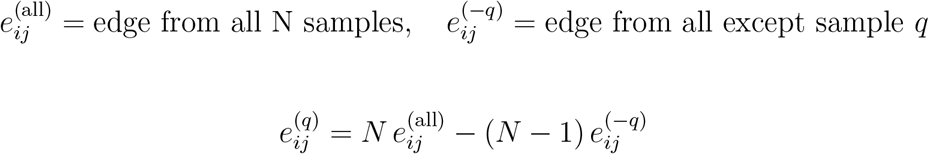

Module scoring is used to measure the strength of correlations between a cluster of gene expressions, using the personalized weights e(q) and the modules in the gene cluster belonging to Ek.

### Module scores

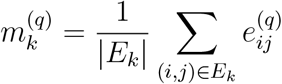

a sigmoidal activation on a linear projection of module score k and learned weight beta of K is then used to predict the basal or classical subtype, which allows the model to determine the most influential genes in a cluster with replication timing. If the probability of the basal subtype is above 50 percent, it is classified as basal.

### Subtype prediction

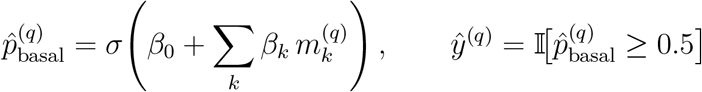

where *σ*(*z*) = 1*/*(1 + *e*^−z^). Any linear or margin-based classifier on 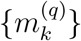 is equivalent in this framework.

Z metric significane is determined using AUC, connectivity, or correlation for x real. mew is using the mean of the metrics for the shuffle modules and sigma is the standard deviation. the empirical p value is calculated using the count of the number of shuffle metrics greater than the real value overr the total number of shuffle iterations. Jaccard stability is computed using the metrics from bootstrap A and B, where E is the expected metric value.

### Null comparison and stability

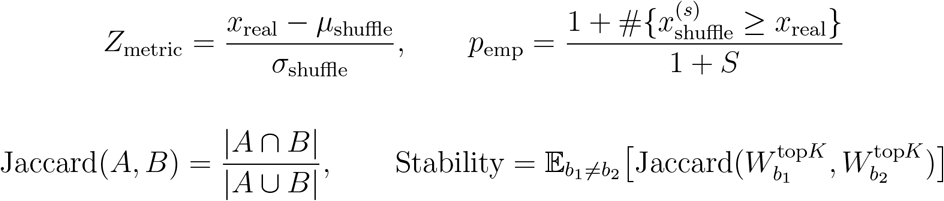

Using the same network construction framework, morphology–RNA networks were generated from 1300-dimensional UNIv2 patch embeddings extracted at 40x magnification using Trident. Patch embeddings for each slide were averaged to obtain a patient-level morphological feature vector. These features were standardized and incorporated into the module-scoring stage of the network pipeline, where morphology was used to weight module activation and influence the gene clusters selected for subtype prediction. Morphology was not used to create separate morphological nodes; rather, the morphological representation modulated the scoring of RNA-derived modules within each patient’s LIONESS network. All embeddings were extracted consistently using the same preprocessing parameters to ensure reproducibility.

## 3 Results

Preprocessing, Construction, and statistical validation were performed on the Ohio State Supercomputer using NVIDIAA100 GPUS. The scripts took 30 minutes to assemble the networks and perform the subtyping.

The number of Early-Early replication timing pairs (EE), Early-Late (EL), and Late-Late gene replication pairs (LL) is measured for each variable. Jaccard stability is shown to be stable due to the small geneset and large number of bootstraps. Density and transitivity plots show the shuffle plots agglomerate, demonstrating variance is reproducible and is not due to dataset artifacts.

**Figure 1.**
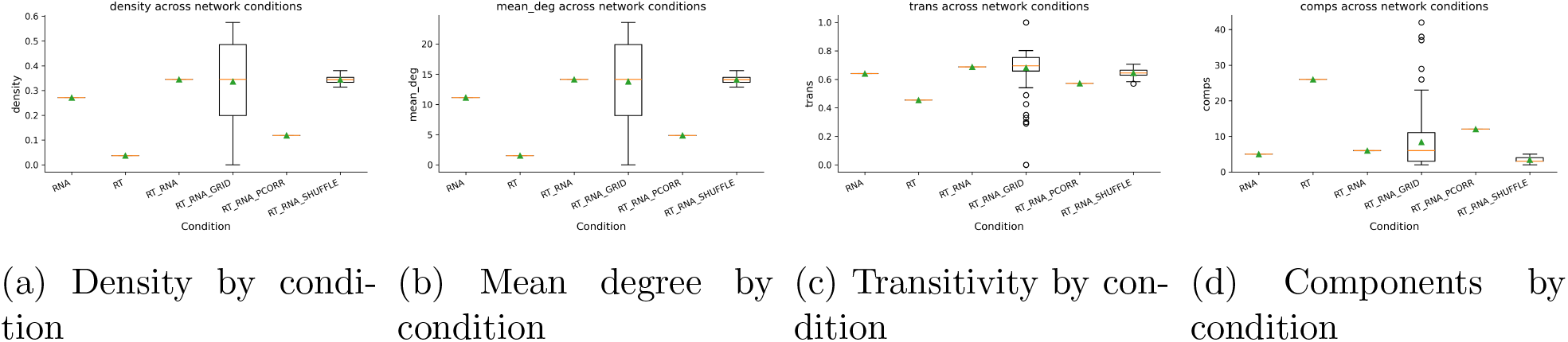
Global network topology across conditions. Boxplots summarize density, mean degree, transitivity, and number of connected components for each folder (e.g., RT, RT_RNA, RT_RNA_PCORR, grid runs, and shuffles).

The results from the tables shown below demonstrate that the individualized network subtyping is not sufficient using the 50 gene systems, with signals barely above noise. RT cannot be modeled as nodes due to RT encompassing multiple genes.

The module level predictions demonstrate that RT and the RT and RNA module have the same performance using the same 17 genes across a single gene cluster module. the peak AUC of .8 across the module level condtions and the increased edge correlation across both implies that the RT addition increases the robustness of the network while maintaining the same performance. The replication timing provides a correlative view of gene coexpression mechanisms, but does not improve the accuracy of the output, as the accuracy depends on the strength of the gene clusters.

The RT and RNA enrichment shows a natural overlap of genesets across the RT and RNA conditions respectively. specifically, the genes used have an 88 percent overlap with the genes used in the other conditions. One of the shuffle graphs for the global networks is shown for the RT and RNA condition as well as a lioness network, where consistent edge networks are shown, with no anomalous of obscure connections.

Gene networks across each case saw higher edge weights with a higher prevalence, validating the evaluation of genetic correlational strength across gene clusters. Across conditions, the RNA grid shifting demonstrated the highest variability, while the control shuffle module generated little shift and stable overlap. Tabke 6 sees that 16 genes used in the module 0 for RT and RNA modules have overlap with the PURIST gene set, which has a reported subtyping AUC of 99 percent. the accuracy of subtyping in this case relies on the strength of gene clusters rather than the global network strength. The distribution of edges is shown in Figure 5, where this is a high count of low weight edges that are filtered from the networks. As expected, the RNA and RT grids and shuffle would see a much higher number of edges than the regulated global networks of RNA +RT. partial correlation was not used, and RT was unable to generate edge weights due to the constraint of having now gene weights.

**Table 2:** Cross-validated subtype classification performance (AUC) across conditions for basal and classical pancreatic ductal adenocarcinoma using individualized networks. Values are taken exactly as provided in the CSV (no column changes). AUC values summarize discrimination between basal and classical subtypes using the specified network inputs. Reported columns are preserved exactly from the source file.

| condition | subtype_auc |
| --- | --- |
| RNA | 0.540608 |
| RT | NaN |
| RT_RNA | 0.532275 |

**Table 3:** Module-level AUCs derived from RNA-only individualized networks. These results show which modules are most informative for subtype prediction. Columns are preserved as in the source file. This table highlights RNA-only network modules contributing to classification; use alongside RT and RT+RNA module tables to compare signal localization.

| module | n_genes | auc |
| --- | --- | --- |
| module_0 | 17 | 0.805688 |
| module_2 | 6 | 0.595767 |
| module_1 | 15 | 0.380820 |

**Table 4:**
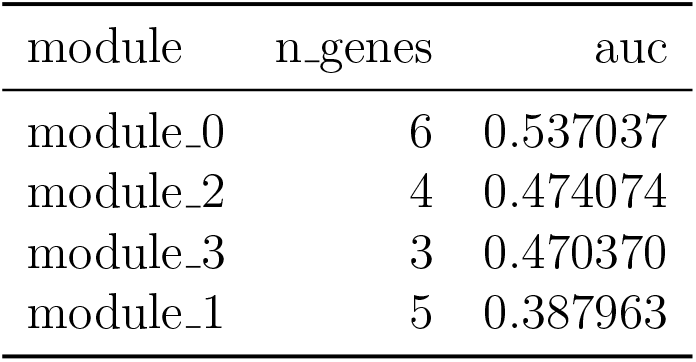
Module-level AUCs derived from replication-timing–proxy features alone. Columns are preserved as in the source file. These results isolate the contribution of replication-timing proxies to subtype prediction independent of RNA.

| module | n_genes | auc |
| --- | --- | --- |
| module_0 | 6 | 0.537037 |
| module_2 | 4 | 0.474074 |
| module_3 | 3 | 0.470370 |
| module_1 | 5 | 0.387963 |

**Table 5:**
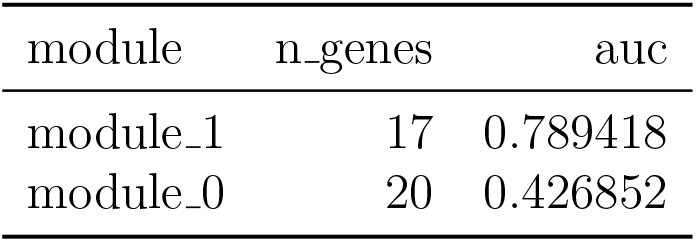
Module-level AUCs from joint replication-timing proxy and RNA networks, representing the integrated model. Columns are preserved exactly as provided. Compare these values with RNA-only and RT-only tables to identify modules where integration improves stability or performance.

| module | n_genes | auc |
| --- | --- | --- |
| module_1 | 17 | 0.789418 |
| module_0 | 20 | 0.426852 |

**Table 6:** Mapped genes for module 1 with HGNC symbols and descriptions (from module_1_genes_mapped.txt).

| Gene | Ensembl ID | Description |
| --- | --- | --- |
| LAMC2 | ENSG00000058085 | laminin subunit gamma 2 [Source:HGNC Symbol;Acc... |
| COL17A1 | ENSG00000065618 | collagen type XVII alpha 1 chain [Source:HGNC S... |
| DHRS9 | ENSG00000073737 | dehydrogenase/reductase 9 [Source:HGNC Symbol;A... |
| SLC2A1 | ENSG00000117394 | solute carrier family 2 member 1 [Source:HGNC S... |
| TNS4 | ENSG00000131746 | tensin 4 [Source:HGNC Symbol;Acc:HGNC:24352] |
| SCEL | ENSG00000136155 | sciellin [Source:HGNC Symbol;Acc:HGNC:10573] |
| TMPRSS4 | ENSG00000137648 | transmembrane serine protease 4 [Source:HGNC Sy... |
| MAL2 | ENSG00000147676 | mal, T cell differentiation protein 2 [Source:H... |
| SPRR3 | ENSG00000163209 | small proline rich protein 3 [Source:HGNC Symbo... |
| SPINT2 | ENSG00000167642 | serine peptidase inhibitor, Kunitz type 2 [Sour... |
| KRT19 | ENSG00000171345 | keratin 19 [Source:HGNC Symbol;Acc:HGNC:6436] |
| CST6 | ENSG00000175315 | cystatin E/M [Source:HGNC Symbol;Acc:HGNC:2478] |
| LEMD1 | ENSG00000186007 | LEM domain containing 1 [Source:HGNC Symbol;Acc... |
| S100A14 | ENSG00000189334 | S100 calcium binding protein A14 [Source:HGNC S... |
| S100A2 | ENSG00000196754 | S100 calcium binding protein A2 [Source:HGNC Sy... |
| LAMB3 | ENSG00000196878 | laminin subunit beta 3 [Source:HGNC Symbol;Acc:... |
| KRT6A | ENSG00000205420 | keratin 6A [Source:HGNC Symbol;Acc:HGNC:6443] |

**Figure 2.**
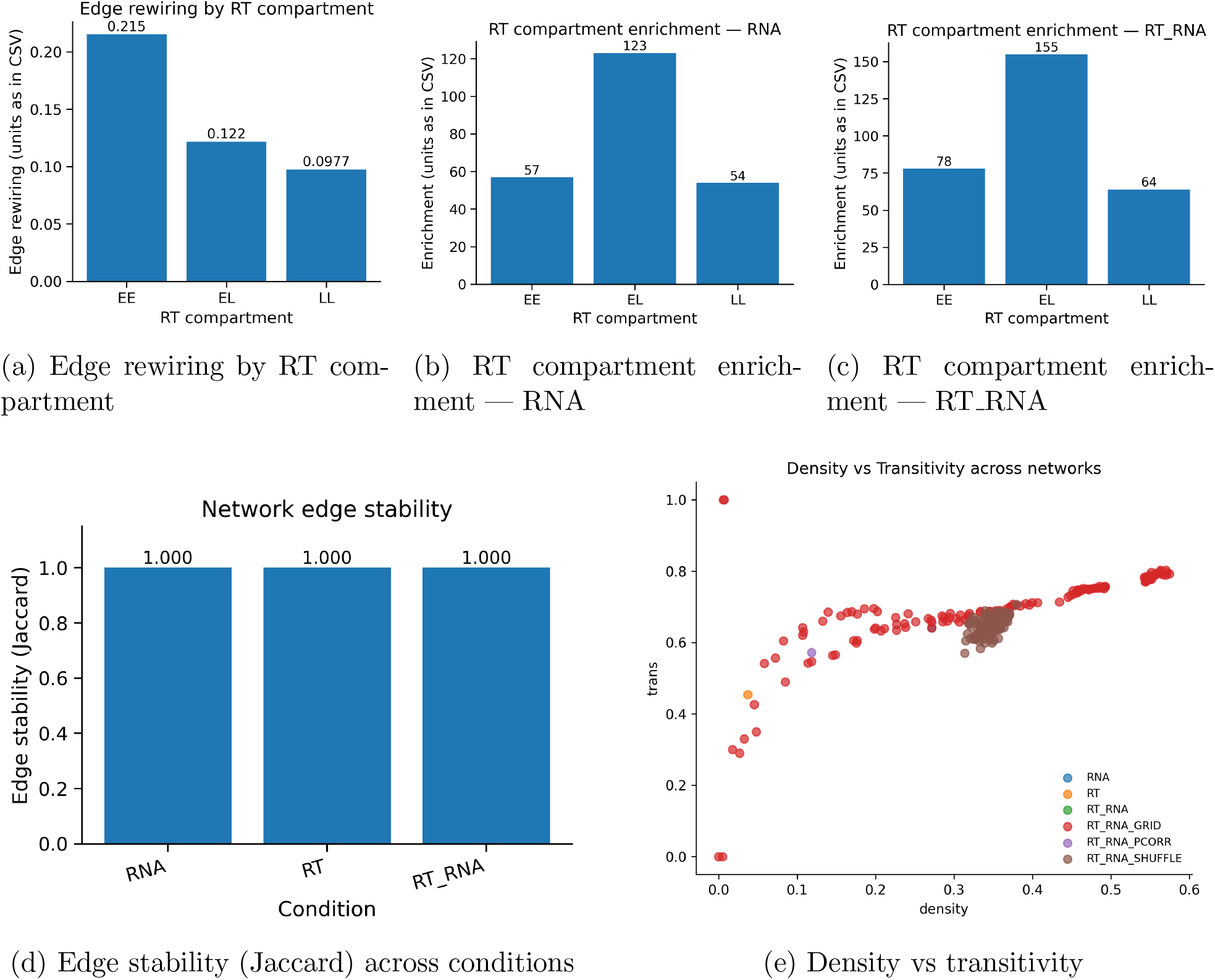
Replication timing context and stability. Top: rewiring and enrichment across early/late compartments for RNA and RT RNA conditions. Bottom-left: edge stability by condition. Bottom-right: relationship between density and transitivity across networks.

**Figure 3.**
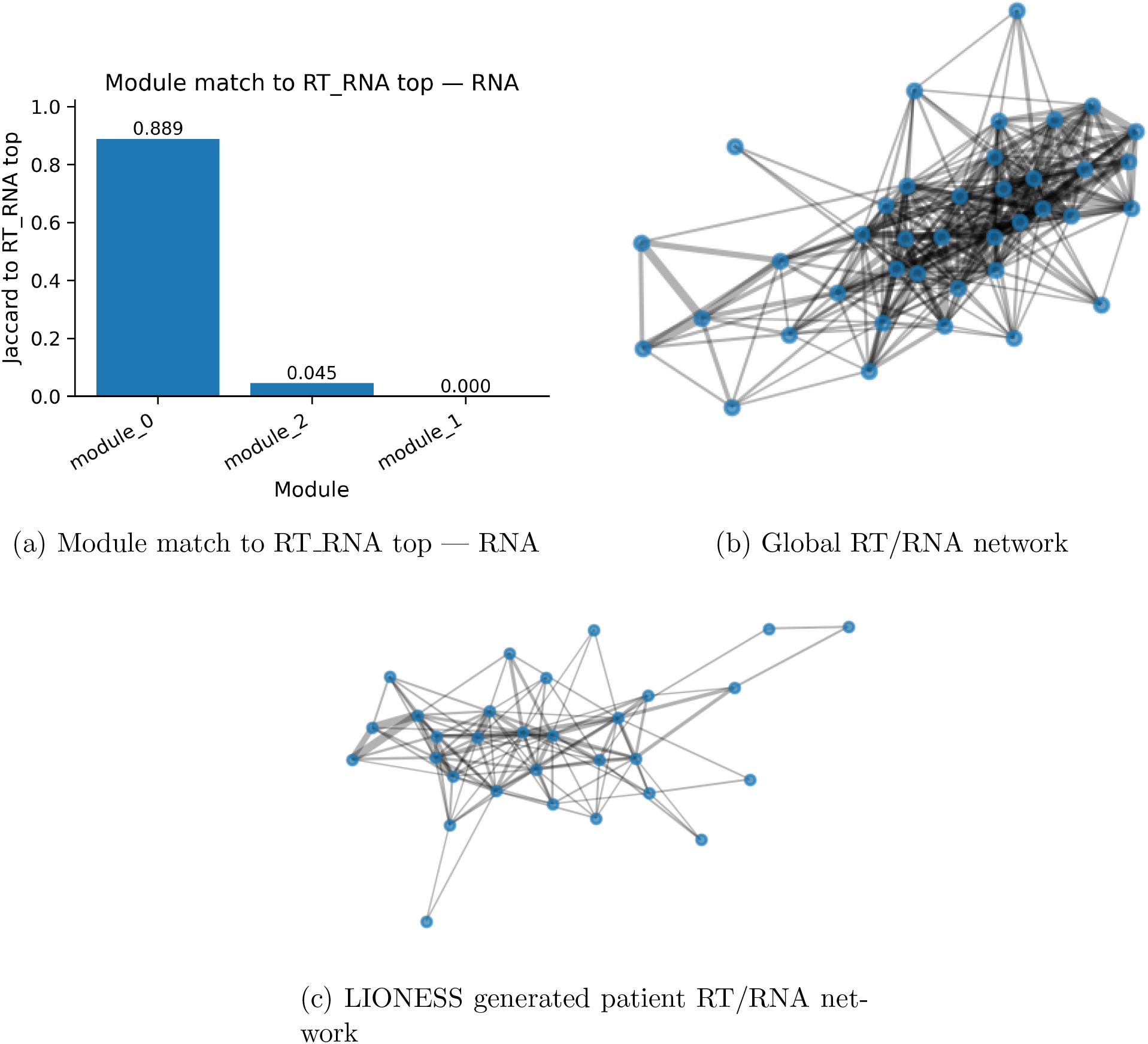
Module-level Jaccard overlap with the RT RNA top modules. Bars show permodule similarity, highlighting modules stabilized or strengthened when replication timing proxies are integrated. Global and personal networks for RT/RNA conditions are shown

**Figure 4.**
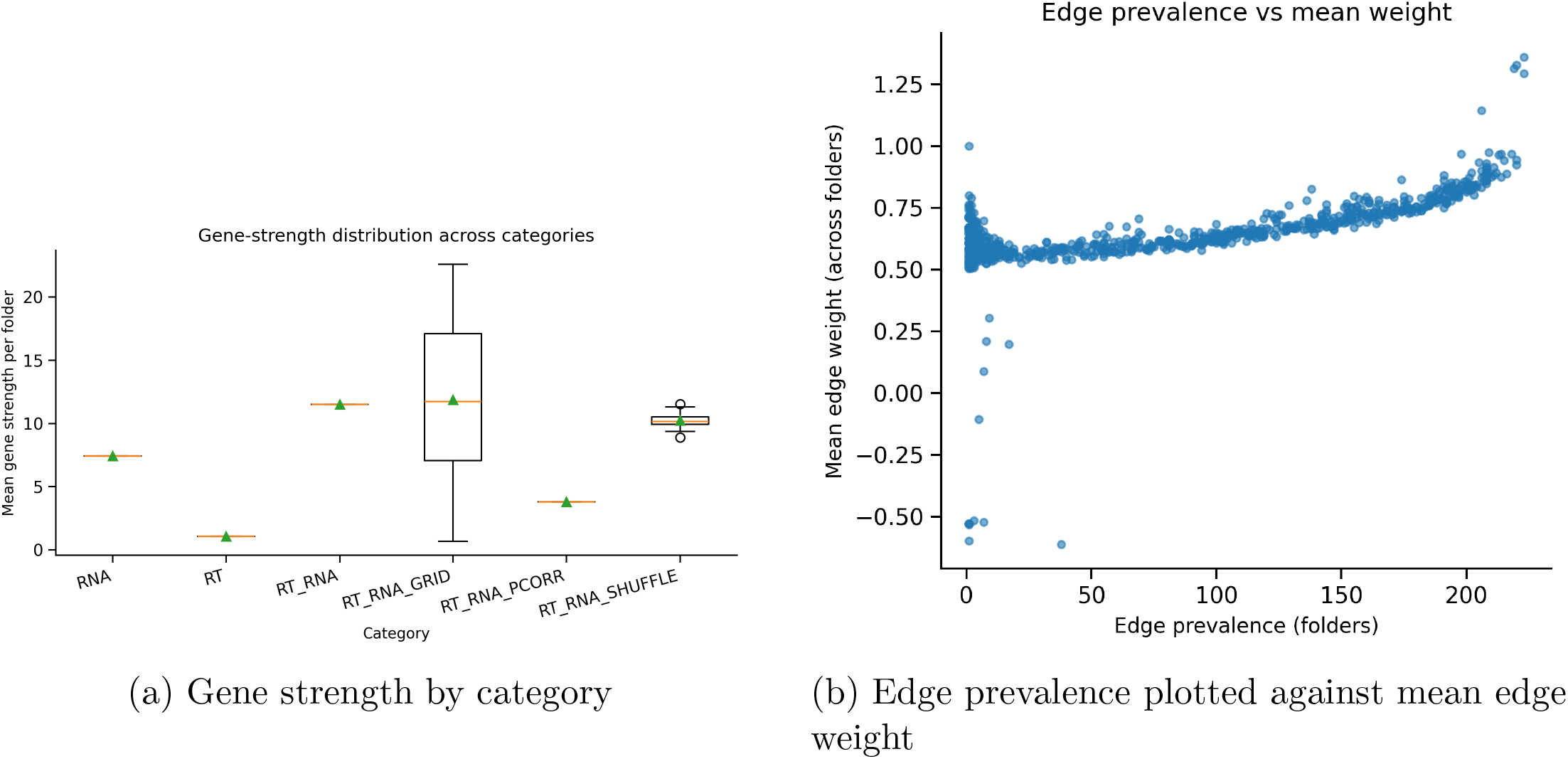
Gene strength by category for each condition, as well as the edge prevalence and the mean weight.

**Figure 5.**
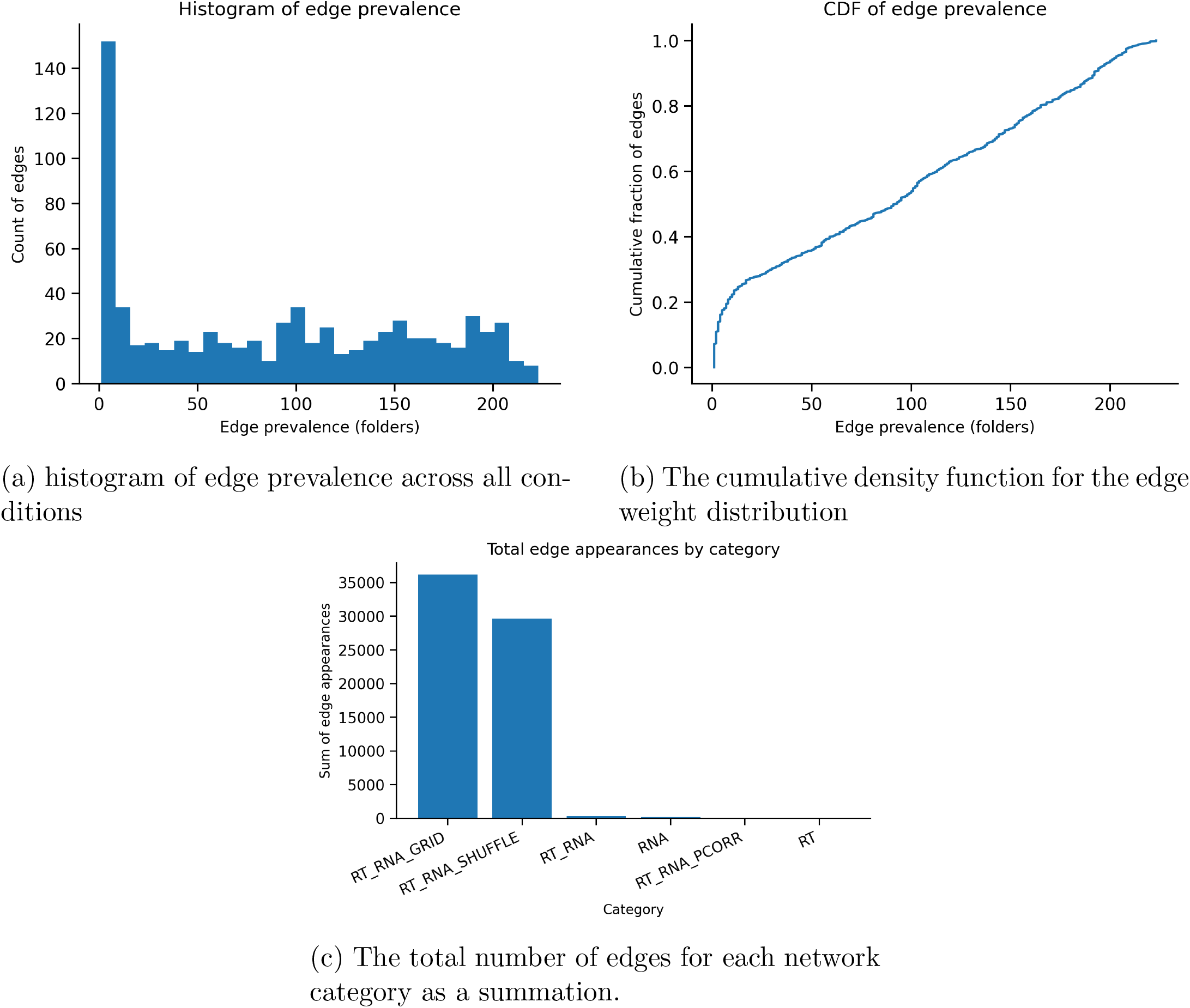
Distribution and frequency of edge weights across network construction conditions. (a) Histogram of edge prevalence showing how often individual edges appear across RNA, RT, grid-search, and shuffle network variants. (b) Cumulative distribution function of edge weights illustrating the overall edge-weight distribution after thresholding and top-K selection. (c) Total number of edge appearances aggregated by network category, highlighting the larger edge counts produced by grid-search and shuffle conditions compared with the constrained global RNA and RT+RNA networks.

Morphology–RNA networks constructed using UNIv2 embeddings were used to influence module scores and edge weights, resulting in a skewed variance distribution across edges. The control-shuffle groups demonstrated that this variance was not due to dataset artifacts, as shown in Figure 6. It is important to note that one global morphology graph and one global RNA graph were constructed, meaning the number of edges across these aggregated cases is negligible compared to the summated edge weights of the shuffle and grid groups. The mean degree was highly variant across the grid condition, as expected, compared to the RNA and morphological networks, and the RNA networks also presented high stability. Edge prevalence as a function of mean weight remained relatively stable, though left-skewed. The appearance of the global morphological–RNA network is shown in Figure 6J. These results demonstrate that morphology-based networks derived from ViT embeddings can be stably correlated with genetic coexpression within the Moffitt gene cohorts [7,10,11,15].

**Figure 6.**
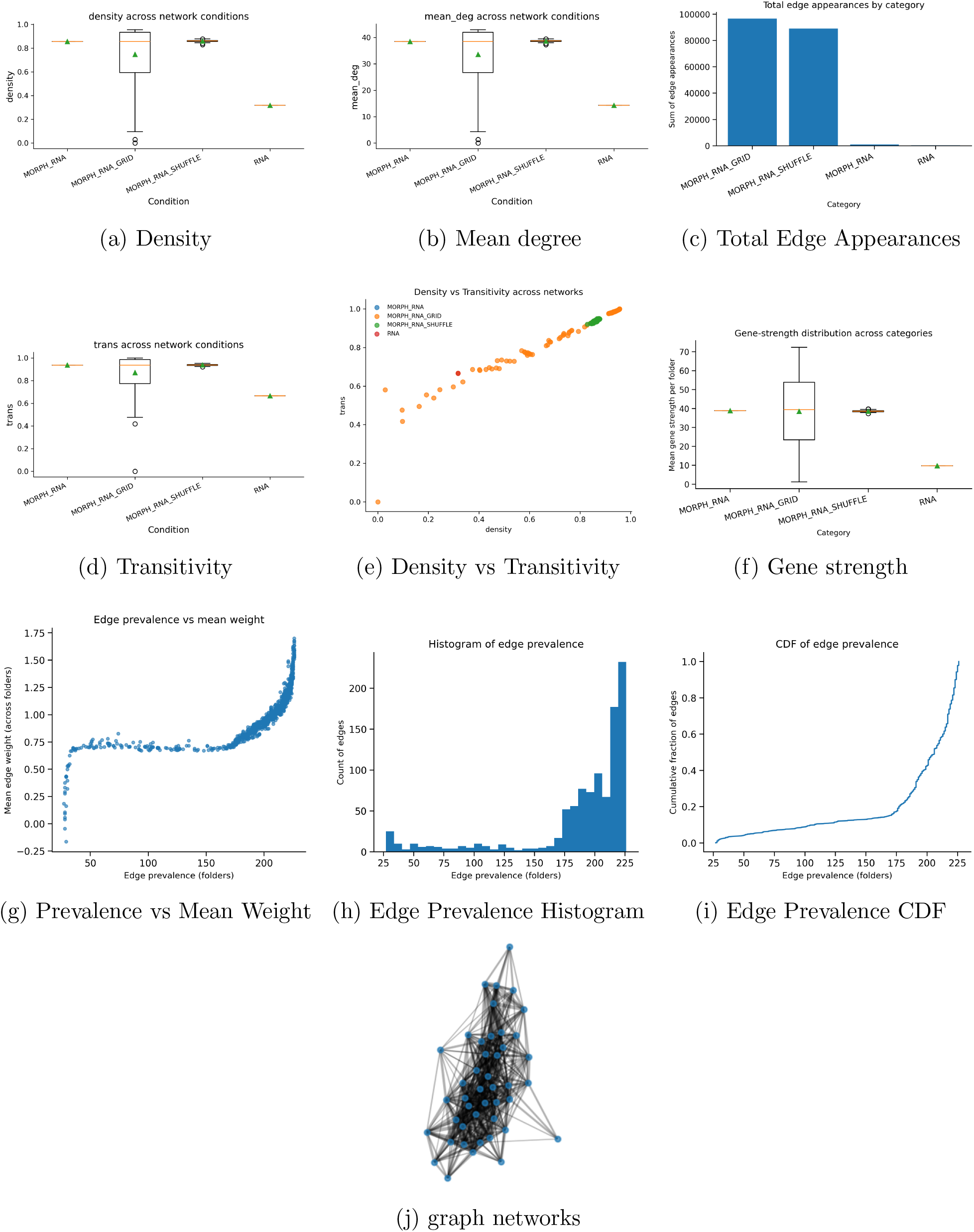
All morphology–RNA network diagnostics. Panels summarize global topology, transitivity, edge prevalence behavior, gene strength metrics, and edge-weight structure across network conditions.

The tables for the morphological lioness networks demonstrated a dipole-like shift, whereby morphological embeddings emphasized one subtype heavily and chose genes corresponding to modules related to the respective subtype. Module 0 shows a high basal subtype preference, with AUC lowered due to ambiguity between low and high confidence samples. The second module selected 19 Classical genes and had a lower AUC than the basal cohort, most likely due to natural distributions of moffitt classifications, by which basal samples are more likely to be high confidences, while most classically subtyped patients align towards low confidence or ambiguous subtypes. Regardless, Using the morphological characteristics of whole slide images, the feature mapping between genes for the classical and basal cohort can be derived. This presents a novel means to wholistically map physical features from pathology images to genetic co-expression.

**Table 7:** Module-level AUCs for Morph RNA LIONESS networks.

| **Module** | **n_genes** | **AUC** |
| --- | --- | --- |
| module_0 | 26 | 0.7496 |
| module_1 | 19 | 0.3294 |

**Table 8:** Subtype classification AUCs for global RNA and Morph RNA network conditions.

| **Condition** | **Subtype AUC** |
| --- | --- |
| RNA | 0.6168 |
| MORPH_RNA | 0.5798 |

**Table 9:** Subtype discrimination of RT RNA network modules using LIONESS-based connectivity. For each module, a patient-specific score is computed as the mean edge weight among all intra-module edges in that patient’s RT RNA_LIONESS network; AUC is calculated with `classical` as the positive class and `basal` as the negative class.

| Module | Number of genes | AUC (classical as positive) | AUC (basal as positive)<sup>a</sup> |
| --- | --- | --- | --- |
| module_1 | 17 | 0.42 | 0.58 |
| module_0 | 20 | 0.37 | 0.63 |

<sup>a</sup>Computed as $1 - \text{AUC}_{\text{classical}}$ to reflect discrimination in the basal direction.

**Table 10:** Distribution of Moffitt subtypes in the TCGA PAAD cohort and its implication for LIONESS-based module AUCs. Because basal cases are more frequent, modules whose connectivity is systematically higher in basal tumours will yield AUC < 0.5 when `classical` is treated as the positive class, even though they show meaningful discrimination in the basal direction.

| Subtype | Number of cases | Proportion of cohort (%) |
| --- | --- | --- |
| Basal | 110 | 60.1 |
| Classical | 73 | 39.9 |

## 4 Discussion

Network biology offers a means to understand gene coexpression given previously understood functions of genes. By examining coexpression and clustering strength, gene expression can sometimes be inferred from stronger or more central genes within a cluster, consistent with network theory and biological graph principles [10,11]. Spatial transcriptomics models and digital pathology approaches typically integrate genes as large feature vectors, which may introduce noise if features are not thresholded or selected appropriately. Methods such as high-variance gene selection do not capture biological interpretability, while ontology-based selection does not always maximize predictive strength. Gene ontology can associate gene expression with morphological patterns across tissue regions, providing a means to infer expression across samples without re-estimating expression from morphological features directly.

Network biology in the replication-timing domain provides additional insight into how gene expression is influenced by chromatin structure and temporal replication programs [1,2,17,18]. The accuracy of these models depends primarily on the strength of correlations among genes rather than the specific numerical metrics used. Genetic networks therefore hold potential for quantifying and correlating expression across biological structures while incorporating the replication-timing dimension [1,2,17].

Although replication timing may not improve subtyping accuracy, it improves robustness by emphasizing genes with strong regulatory signals. Replication timing has also been shown to stabilize network properties such as density and transitivity, consistent with long-range epigenetic organization and replication-domain structure [3,4]. This suggests that gene expression can be modeled with respect to chromatin stability and genome-regulatory architecture.

Morphological embeddings also demonstrated strong capacity to detect and score modules corresponding to Basal-Like and Classical subtypes, aligning with previous findings on PDAC subtype structure and morphology-linked differentiation states [15,16]. This supports modeling morphology alongside gene expression. Future biological networks may integrate protein cascades and morphological networks, while chromatin-based features and mutations operate as “invisible variables” that indirectly influence cascade behavior.

In digital pathology, multimodal models are increasingly necessary to annotate slides with biological mechanisms not directly visible in raw histology [1,2,7]. By connecting morphological features to gene expression, models can learn to infer biological cascades based on variations in cell neighborhoods, cell phenotypes, and patch-level motifs.

## 5 Limitations

Partial correlation networks were not measured at the patient level, but LIONESS networks were constructed [7]. The AUC is not strictly dependent on edge strength, but on which genes are included within modules influenced by morphological inputs. Increasing the number of nodes or genes lowers network resolution. STRING database connections were attempted; however, connections among many Moffitt genes were incomplete or absent, which is expected for sparse or poorly annotated regulatory interactions [8,9]. The globally constructed networks are not subtype-specific at the patient level but can characterize cohort-level variance related to tumor progression. Nodes were not constructed from morphology and replication timing directly; instead, these modalities influenced edge weights and module scoring.

## 6 Conclusions

Personalized replication-timing–based correlative networks provide insight into how chromatin structure shapes gene expression variation across patients [1,2,17,18]. This can help characterize cohort-level differences between Classical and Basal-Like populations [15,16]. Although replication timing did not improve predictive performance, it offers an effective means to approximate chromatin stability at the case level. Morphologically influenced networks demonstrate that image embeddings and phenotypes connect meaningfully to PDAC subtype structure [15,16]. Vision transformer embeddings and other image-derived features can therefore be used to understand gene coexpression directly. Future work includes modeling downstream molecular consequences of gene expression, including protein-cascade signaling, in relation to morphological features.

## 7 Ethics Statement

All datasets are public and anonymized.

## 8 Author Contributions

All work was performed by the first author, with supervision by the second.

## 10 Data Availability

TCGA PAAD is a publically available pancreatic cancer dataset. All code is available in the repository: https://github.com/Alejandro21236/LIONESS4PATH

## References

[1] Hansen, R. S., Thomas, S., Sandstrom, R., Canfield, T. K., Thurman, R. E., Weaver, M., Dorschner, M. O., Gartler, S. M., & Stamatoyannopoulos, J. A. “Sequencing Newly Replicated DNA Reveals Widespread Plasticity in Human Replication Timing.” Proceedings of the National Academy of Sciences, vol. 107, no. 1, 2010, pp. 139–144. 10.1073/pnas.0912402107.

[2] Rivera-Mulia, J. C., & Gilbert, D. M. “Replication Timing and Transcriptional Control: Beyond Cause and Effect—Part II.” Current Opinion in Genetics & Development, vol. 37, 2016, pp. 106–113. 10.1016/j.gde.2016.03.002.

[3] Pope, B. D., Ryba, T., Dileep, V., Yue, F., Wu, W., Denas, O., Vera, D. L., Wang, Y., Hansen, R. S., Canfield, T. K., Thurman, R. E., Weaver, M., Sandstrom, R., Reynolds, A. P., Thurman, A. L., Stamatoyannopoulos, J. A., Collins, F. S., Crawford, G. E., Dekker, J., & Gilbert, D. M. “Topologically Associating Domains Are Stable Units of Replication-Timing Regulation.” Nature, vol. 515, no. 7527, 2014, pp. 402–406. 10.1038/nature13986.

[4] Fortin, J. P., & Hansen, K. D. “Reconstructing A/B Compartments as Revealed by Hi-C Using Long-Range Correlations in Epigenetic Data.” Genome Biology, vol. 16, 2015, p. 180. 10.1186/s13059-015-0741-y.

[5] Zhou, W., Triche, T. J., Laird, P. W., & Shen, H. “SeSAMe: Reducing Artifactual Detection of DNA Methylation by Infinium BeadChips in Genomic Deletions.” Nucleic Acids Research, vol. 46, no. 20, 2018, e123. 10.1093/nar/gky691.

[6] Shetty, A. V., et al. “Single Cell Multiomics Approach to Analyze Replication Timing and Gene Expression in Mouse Preimplantation Embryos.” Communications Biology, vol. 8, no. 1, 2025. 10.1038/s42003-025-08694-5.

[7] Kuijjer, M. L., Tung, M. G., Yuan, G., Quackenbush, J., & Glass, K. “Estimating Sample-Specific Regulatory Networks.” iScience, vol. 14, 2019, pp. 226–240. 10.1016/j.isci.2019.03.001.

[8] Marbach, D., Costello, J. C., Küffner, R., Vega, N. M., Prill, R. J., Camacho, D. M., Allison, K. R., Kellis, M., Collins, J. J., & Stolovitzky, G. “Wisdom of Crowds for Robust Gene Network Inference.” Nature Methods, vol. 9, no. 8, 2012, pp. 796–804. 10.1038/nmeth.2016.

[9] Glass, K., Huttenhower, C., Quackenbush, J., & Yuan, G. “Passing Messages Between Biological Networks to Refine Predicted Interactions.” PLoS ONE, vol. 8, no. 5, 2013, e64832. 10.1371/journal.pone.0064832.

[10] Newman, M. E. J. Networks: An Introduction. Oxford University Press, 2010.

[11] Watts, D. J., & Strogatz, S. H. “Collective Dynamics of ‘Small-World’ Networks.”Nature, vol. 393, no. 6684, 1998, pp. 440–442. 10.1038/30918.

[12] Jaccard, P. “Étude Comparative de la Distribution Florale dans une Portion des Alpes et des Jura.” Bulletin de la Société Vaudoise des Sciences Naturelles, vol. 37, 1901, pp. 547–579.

[13] Good, P. Permutation, Parametric, and Bootstrap Tests of Hypotheses. 3rd ed., Springer, 2005.

[14] Efron, B., & Tibshirani, R. J. An Introduction to the Bootstrap. CRC Press, 1994.

[15] Moffitt, R. A., Marayati, R., Flate, E. L., Volmar, K. E., Loeza, S. G. H., Hoadley, K. A., Rashid, N. U., Williams, L. A., Eaton, S. C., Chung, A. H., Smyla, J. K., Anderson, J. M., Kim, H. J., Bentrem, D. J., Talamonti, M. S., Iacobuzio-Donahue, C. A., Hollingsworth, M. A., & Yeh, J. J. “Virtual Microdissection Identifies Distinct Tumor- and Stroma-Specific Subtypes of Pancreatic Ductal Adenocarcinoma.” Nature Genetics, vol. 47, no. 10, 2015, pp. 1168–1178. 10.1038/ng.3398.

[16] Rashid, N. U., Peng, X. L., Jin, C., Moffitt, R. A., Volmar, K. E., Belt, B. A., Panni, R. Z., Mehta, A., Lee, S. M., Hurt, C., Manjunath, Y., & Yeh, J. J. “PURIST: Pancreatic Cancer Uniform Classification by Transcriptomics.” Clinical Cancer Research, vol. 26, no. 16, 2020, pp. 4001–4009. 10.1158/1078-0432.CCR-19-3784.

[17] Mukherjee, S., & Callegari, A. J. “How Replication Timing Shapes Genome Maintenance and Cancer Genome Evolution.” Frontiers in Cell and Developmental Biology, vol. 9, 2021, 671120. 10.3389/fcell.2021.671120.

[18] Rhind, N., & Gilbert, D. M. “DNA Replication Timing.” Cold Spring Harbor Perspectives in Biology, vol. 5, no. 8, 2013, a010132. 10.1101/cshperspect.a010132.

